# Principles of organelle membrane bridging established using cytosolic tether mimics

**DOI:** 10.1101/2020.09.02.279398

**Authors:** Mohammad Arif Kamal, Josip Augustin Janeš, Long Li, Franck Thibaudau, Ana-Suncana Smith, Kheya Sengupta

## Abstract

The interactions between different intra-cellular organelles, including the endoplasmic reticulum, have recently been in focus thanks to the tremendous progress in imaging them using cryogenic transmission electron microscopy. However, they are still difficult to study in cellulo, and reconstituting these systems has been a standing challenge. Here we achieve this task using a giant unilamellar vesicle (GUV) and supported lipid bilayer (SLB) system. The tethers, which may reside in the cytosol when unbound, are mimicked by single (or double) stranded DNA sequences of two different lengths with ends that are self-sticky, and with terminal cholesterol moieties which insert into GUV or SLB membranes. The DNA-tethers, bound by their sticky-end, can exist in two possible states - either with both cholesterols in the same membrane or each cholesterol in a different membrane, the latter conformation leading to adhesion. Exchange of tether-molecules between the membranes occurs through the aqueous phase. By developing theoretical arguments that are supported in our experiments, we show that this possibility of exchange and the relative difference in the projected area between the two states drives the adhesion due to collective entropic considerations, rather than the usually considered enthalpy of binding. The establishment of this fundamentally different interaction between two membranes suggests that in physiological conditions, the regulation of contact formation inside cells may be very different from the case of the much studied ligand-receptor pairing on the external cell membrane.

## Introduction

The correct organization of organelles, essential for the coherent functioning of eukaryotic cells, is maintained via specialised patches called organelle-contacts [6]. While early electron microscopy images revealed the existence of these patches, their dynamical nature, molecular machinery and function is only beginni,ng to be explored [6, 13, 17, 20, 24, 29]. It is now clear that these contacts not only provide structural scaffolding but also act as important sites of inter-organelle communication, exchanging proteins, small molecules, and lipids across the organelle membranes [7]. These contact sites are dynamic in size and can change in response to environmental or cellular stimuli; however, their regulation is still not established. [7] Nonetheless, it is increasingly clear these structurally-confined contacts between membranes are mediated by anchoring molecules, somewhat reminiscent of cell surface ligand-receptor binding that mediates cell adhesion. In analogy with adhesion receptors that are sometimes called “linkers”, the proteins that span the inter-organelle gap and physically keep two organelles together are termed “tethers” [6]. It is now becoming clear that many, possibly all, organelles communicate by means of such inter-organellar tethering. [6, 7]

In spite of recent progress, very little is known about the nature of the contact sites and the molecules that mediate their formation. Two broad groups of tethers have been identified, with very different length and flexibility [5]. Some of these tethers reside on the membrane, whereas some others are soluble cytosolic molecules that are recruited to the membranes dynamically. Many are expected to undergo major conformation changes upon tethering. While molecular details are definitely required to get a detailed understanding of function, consideration of general thermodynamic and geometric arguments can already help us predict the possibilities of a system, without need of specific chemical details.

In the past, such a generic approach has elucidated a number of physical determinants of membrane anchoring [9, 10, 27, 28]. Particular success was achieved using the paradigm of adhesion between giant unilammelar vesicles (GUVs) and solid supported lipid bilayers (SLBs) [10, 23]. In addition to macro-regulators like the density of ligands and receptors, and the mechanical properties of the membranes [32], more subtle effects like positional entropy of the tethers [9], the presence of repelers [15], jamming related size effects [8], relative tether lengths in a dual-tether system [25] etc. have been shown to play a role. In many of these studies, purified adhesion proteins were used, whereas in some others, model proteins served the purpose.

Purification of intra-cellular tethers remains extremely challenging, not least because very little is known about them. Attachment of trafficking endo and exocytotic vesicles to the ER and Golgi are special cases of inter-organellar contacts, which has been better studied. The focus however, has been on fusion and fission of the membrane [2, 5, 31, 36]. To avoid the bottleneck created by protein purification challenges, artificial proteins, macromolecules or even nucleic acids have been used to mimic tethers and anchors, lipid-grafted DNA tethers used as mimic of the SNARE complex being an early example [4, 37].

Simple synthesis and variability of design make DNA tethers particularly attractive artificial tethers. Besides the capacity to tune the binding by adjusting the length of the sequence used for sense/antisense recognition, their design allows for the systematic modifications of length and flexibility [4]. In addition to being used for tethering [4, 37], DNA constructs were also extensively used as glycocalyx-mimetic spacers [12, 16, 18]. More recently, they have been used as force sensors in hybrid cell-surface systems [3, 38]. DNA-tethers have in fact been used in a plethora of situations including building foam-like structures [11], self-assembly of soft Brownian objects [21], or for exploring thermal control of particle-assemblies [30].

Here we use DNA technology to design mimics of inter-organellar tethers that generate contact between two synthetic membranes of a GUV/SLB system. We use single stranded DNA (ss-DNA) or double stranded DNA (ds-DNA) oligomers, of two different lengths to bridge the membranes and create stable contact patches. In each case, the DNA-tether can insert into a membrane with the help of a cholesterol moiety, and can bind a self-similar DNA-tether, which may itself be inserted into a membrane via a sticky-end. From energetic considerations, we expect that the tethers always dimerise via their sticky-ends, and may exist in three possible states - in solution, in a *cis*-configuration with both cholesterols in the same GUV or SLB membrane forming a ‘U’-shape, and in a *trans*-configuration with one cholesterol in the SLB and the other in a GUV forming a ‘I’-shape. Only the latter *trans*-configuration state can bridge the membranes and create intermembrane contacts; thus, contact formation is accompanied by tether conformation changing from ‘U’-like to ‘I’-like, each with very different effective size, or by the recruitment from the solution into the *trans*-configuration. This leads to a novel adhesion mechanism that is not based on the single-tether sense/antisense recognition affinity, but on the interplay between the soluble pool and the *cis*/*trans* configurations with different geometries. We study tethers based on ss-DNA being highly flexible and behaving as an entropic spring, as well as ds-DNA, which is relatively stiff but contains soft joints. Using their long and short versions we were able to explore the effect of both tether length and flexibility on contact formation in this system where a pool of cytosolic tethers self-regulate contact formation *via* geometric effects.

### DNA tether design and properties

#### Tether structure

We first prepare single stranded DNA (ss-DNA) sequences of *N* =28 (short) or *N* =40 (long) base pairs, out of which 10 base pairs build a sticky end (red colored sequence in Fig. 1b), and the reminder form the backbone (green sequence) attached to a cholesterol moiety *via* a flexible TEG-tether linked to a short single-stranded sequence (black TEG and blue ss-sequence). Two identical segments are allowed to recombine at the sticky ends to make a strand twice as long with a cholesterol at each end. Consequently we obtain two constructs which we denote as short and long ss-tethers, respectively. To create ds-tethers, the short or long ss-DNA sequences are first incubated with appropriate backbone sequences of 12 or 24 base pairs such that they combine using sense/antisense recognition (brown sequences in Figs. 1b). Due to the high hybridization free energy (of the order of 20 *k*_*B*_*T* s where *k*_*B*_ is the Boltzmann constant, and *T* temperature), we assume that all ss-tethers recombine into ds-tethers already in solution. In the final step, two identical ds-sequences also recombine at the sticky ends to make short and long ds-tethers, with a cholesterol at each end.

**Figure 1.**
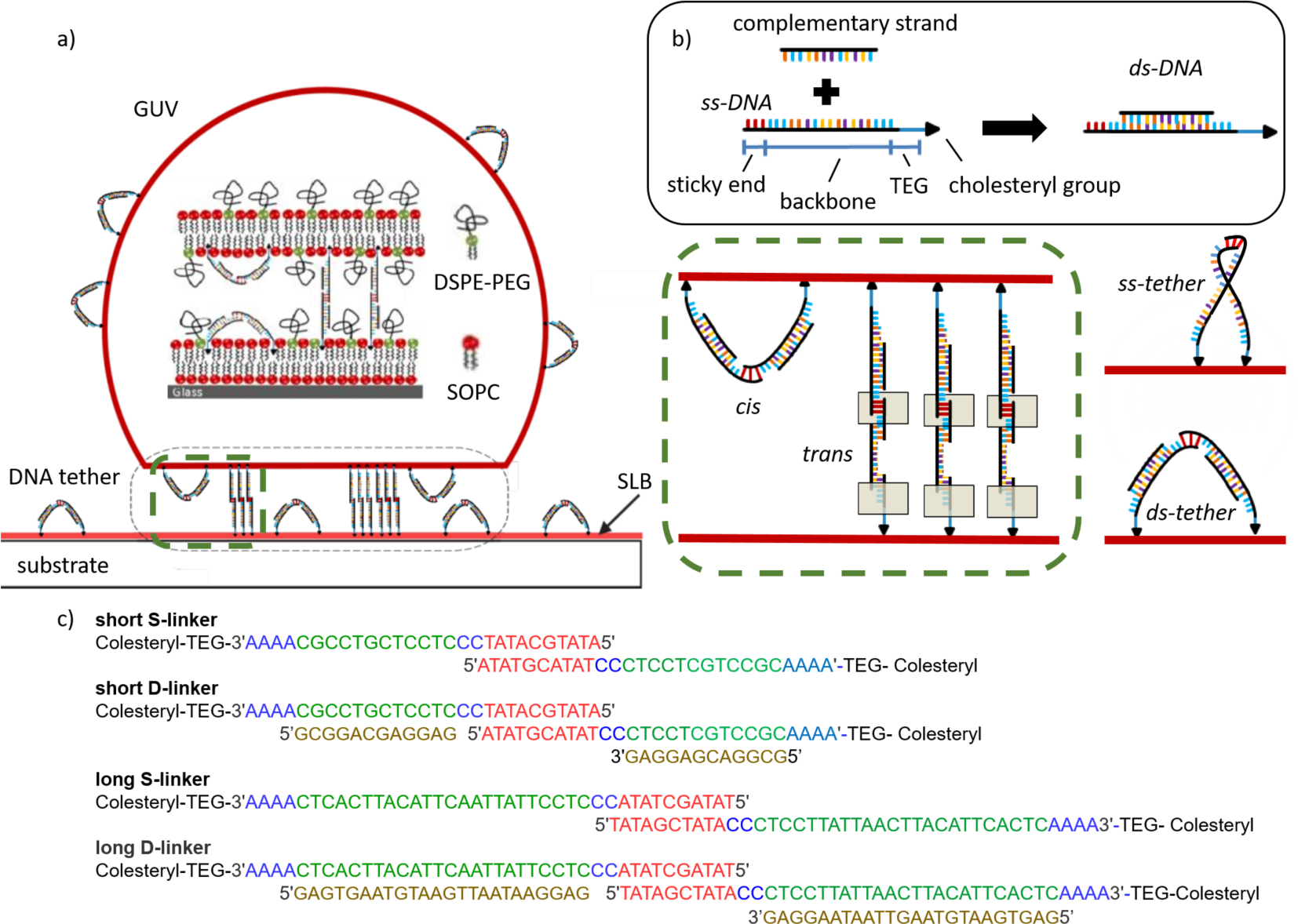
a) Sketch of the GUV-SLB adhesion mediated by DNA tethers. DNA tethers can be either in the non-tethering *cis* (‘U’-shaped) configuration, or in the tethering *trans* (‘I’-shaped) configuration (zoom-in within the green dashed frame). GUV-SLB adhesion is mediated by *trans* tethers. b) Microstructure of the DNA tethers. ss-tethers are made when two identical ss-DNA segments are allowed to recombine at the sticky ends to make a strand twice as long with a cholesterol at each end. ds-tethers are made when two identical ds-DNA segments are allowed to recombine at the sticky ends to make a strand twice as long with a cholesterol at each end. c) Molecular structure, together with the DNA sequences, of the used tether types. TEG stands for Tetra Ethylene Glycol, which links the DNA to the Cholesterol.

In terms of structure, a ds-tether, in its dimerized form, can be considered as a joined rod with three parts: the middle, corresponding to the hybridised sticky-end of 10 base-pairs, is 3 nm long and the two back-bones terminating in the TEG-tag are 4 or 8 nm respectively (calculated using 0.3 nm per base pair). Thus, ignoring the highly flexible TEG and single stranded DNA parts, the length of the long and short tethers in dimerized and extended configuration (cholesterol excluded) is 11 and 19 nm respectively. In terms of flexibility, the ss-tether, being a single stranded DNA with expected persistence length of about 1 nm, is expected to behave as a flexible spring. The ds-tether, on the other hand, is composed of flexibly jointed rigid rods, which allows for a full extension of the structure (Fig. 1).

#### Tether adsorption affinity

In solution, in presence of a lipid membrane, the DNA-tethers are expected to be inserted into the membrane by their cholesterol moiety. This absorption was quantified in by adding fluorescently labeled tethers to a chamber containing a glass supported lipid bilayer (SLB). Adsorption affinity of the tethers was inferred from their SLB-surface densities *ρ*, as quantified using confocal fluorescent microscopy for different bulk concentrations *c*. The relation between density and affinity is given by the Langmuir adsorption isotherm

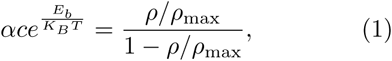

where *K*_*B*_ is the Boltzmann constant, *T* the temperature, *α* a normalization constant and *ρ*_max_ the maximal surface density of tethers. The tether surface density *ρ* is obtained by comparing the fluorescence intensities of the molecules on the SLB with the intensities in the bulk solution (at known bulk concentration *c*). Fitting the measured data to the inverted eq. 1

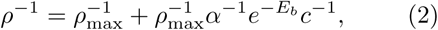

gives the adsorption free energy *E*_*b*_ = 12.4 *k*_*B*_*T*, and *α* = 55.5∗10^−6^(*µM*)^−1^ with *ρ*_max_ = 1.33∗10^12^ cm^−2^ (Fig. 2a). *E*_*b*_ agrees reasonably well with the free energy of cholesterol insertion (9 *k*_*B*_*T*). The discrepancy most likely emerges from the fact that not only the cholesterol, but also the TEG-linker and part of the adjacent single stranded DNA with hydrophobic bases may be inserted. It is nevertheless reasonable to assume that all adsorbed tethers, irrespective of their type, insert both of their cholesterols into the membranes. However, there remains a population of non-absorbed tethers in solution as evidenced by the fact that newly introduced giant unilamellar vesicles (GUVs) become decorated with DNA-tethers even before they approach the SLB, as revealed by transfer of fluorescence to the GUV (Fig 2).

**Figure 2.**
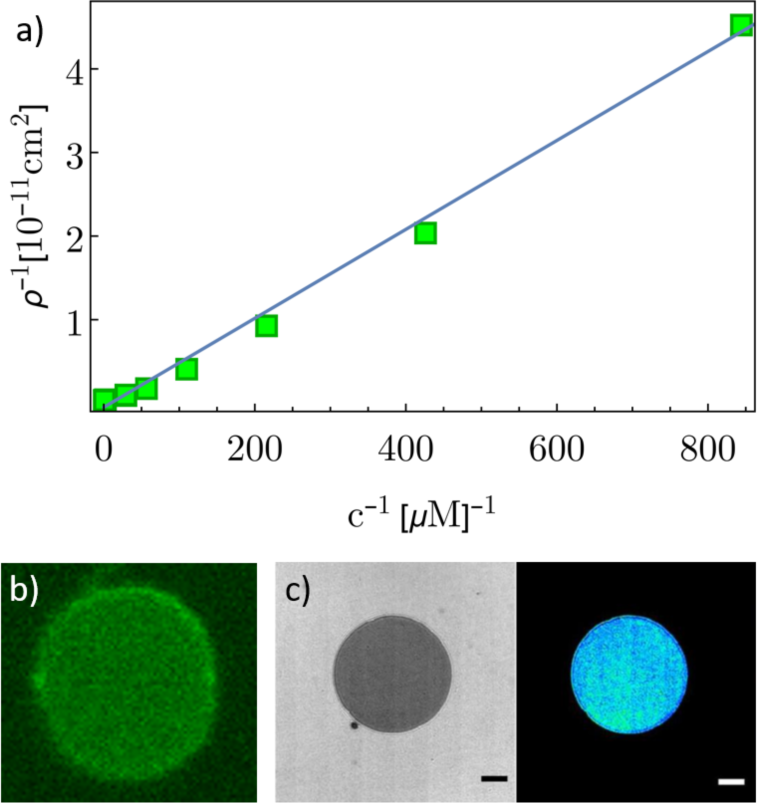
a) Dots correspond to the measured (inverse of the) adsorbed surface density *ρ* for different (inverse of the) bulk concentrations *c* of ds-tethers. The line is the linear fit to eq. 2 with *E*_*b*_ = 12.4 *k*_*B*_*T*. b) Fluorescence image of a free floating GUV, indicating DNA tether adsorption onto the GUV membrane. c) RICM and roughness images of a GUV adsorbed on the SLB.

### Mathematical model of contact regulation

To understand how such soluble tethers may support contact formation, we adapt a statistical mechanical model in which the enthalpy of binding is balanced by total entropy of the system [33, 35]. We model the GUV and the SLB as two equivalent lattices with *N* lattice sites, of which *N*_cz_ < *N* are in a contact-zone, where tethers can mediate the formation of intimate contact - which will be called adhesion domain.

Adsorbed tethers inside the contact-zone can be in one of the two states: (1) *trans*-state, forming an ‘I’-shape, and occupying both a single GUV-lattice site *and* a single SLB-lattice site; or (2) in the *cis*-state, forming a ‘U’-shape, and occupying *ϵ* GUV-lattice sites *or ϵ* SLB-lattice sites. The ‘shape factor’ *ϵ* accounts for the expected difference in the projected area of the *cis*- and the *trans*. Clearly, therefore, *ϵ* ≥ 1, or in other words, the projected area of the *cis*-state on the membrane-plane is assumed to be *ϵ* times larger than that of the *trans* state, and at *ϵ* = 2 the footprint of the *cis* configuration is twice that of the membrane spanning *trans*. Tethers adsorbed outside the contact zone are of course, necessarily in the *cis*-state (Fig.1a).

To keep track of tether numbers in each state, we introduce the following variables; 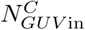 and 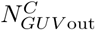, denoting the number of *cis*-tethers inside and outside the vesicle contact zone, 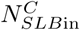 and 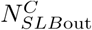, corresponding to the number of *cis*-tethers inside and outside the contact zone on the SLB, and *N*^*T*^ representing the number of *trans*-tethers. The bulk solution surrounding the GUV-SLB system is modeled as an infinite reservoir, characterized by temperature *T* and chemical potential *µ*(*c*), related to the concentration *c* of tethers in the bulk solution. Assuming tethers in both configurations exchange between the membranes and the reservoir, we model the GUV-SLB system in the grand canonical ensemble, with its state function being the grand potential Ω, given by

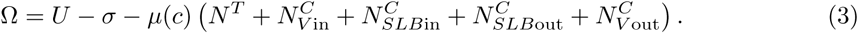

Here *U* is the total binding entalphy of all tethers and *σ* is their entropy of mixing, expressed in terms of *k*_*B*_*T* (*k*_*B*_ being the Boltzmann constant). The last set of terms, proportional to *µ*(*c*), are associated with the free energy change for bringing the tether from the bulk onto the designated part of the GUV or SLB surface. Negative *µ* corresponds to energy cost for absorbing bulk tethers onto the GUV-SLB system, which is decreased by increasing the bulk concentration of tethers (and leading to less negative *µ*).

The enthalpy of the system is given by

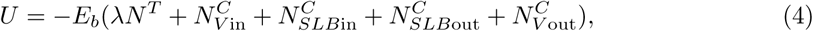

where *E*_*b*_ and *λE*_*b*_ are respectively the *cis*- and *trans*-adsorption energies, with *λ* being a dimensionless number. In the current system*E*_*b*_ = 12.4*k*_*B*_*T*, while it is anticipated that *λ* < 1, making the *cis* binding more likely than the adhesive *trans* state. The reason for this difference is the small free energy cost of transitioning from the *cis*-to the *trans*-state, due to the relative restriction of the internal configurational space of tethers in the *trans*-state. The fact that forming an adhesive contact actually costs free energy makes the tether-binding fundamentally different than the typical ligand-receptor pairing [33, 34].

Total positional entropy is calculated by counting the possible configurations of tethers on the GUV and the SLB, giving:

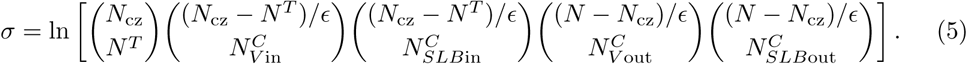

Here we take into account the entropy of *trans* tethers (first term), and *cis*-tethers inside the contact zone on the vesicle (second term) and the SLB (third term), together with the entropy of *cis*-tethers on the vesicle (fourth) and the SLB (firth term) outside the contact zone. Here again, the shape-factor *ϵ* marks an importance difference compared to ligand-receptor binding [33, 34].

Equilibrium density of *trans*-states is calculated by minimizing eq. 3 (see Appendix), with respect to the tether numbers on the SLB and the vesicle in and out of the contact zone, under the constraint of the constant chemical potential [33, 34]. Notably, the concentration of *cis*-tethers on the extended surfaces of the GUV and SLB are related to *µ* by the Langmuir absorption isotherm (Appendix eq. 10). The solution of this minimization is furthermore the density of trans bonds creating a close contact between the two membranes *N*^*T*^ */N*_cz_ (Appendix eq. 8), henceforth called *trans* density (equivalent to the density of ligand-receptor bonds in a classical adhesion system). We explore it as a function of the chemical potential *µ* for different shape and energy factors *ϵ* and *λ* (Fig. 3a-b), to give insight into the mechanisms of contact formation.

**Figure 3.**
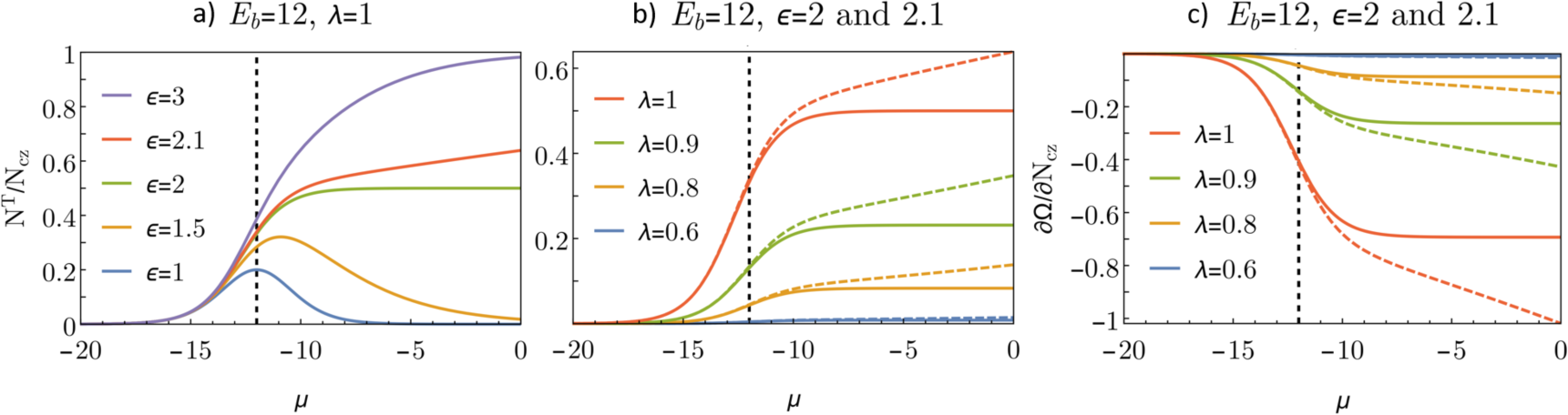
a) Equilibrium density of *trans*-states as a function of the chemical potential *µ* for *λ* = 1 and several values of *ϵ*. b) Equilibrium density of *trans*-states as a function of the chemical potential *µ* for *ϵ* = 2 (full lines) and *ϵ* = 2.1 (dashed lines) and several values of *λ*. c) Spreading pressure ∂Ω*/*∂*N*_cz_ as a function of the chemical potential *µ* for *ϵ* = 2 (full lines) and *E* = 2.1 (dashed lines) and several values of *λ*. Vertical dashed lines are denoting *µ* = −*E*_*b*_ = −12.

We start with the case of identical adsorption affinities of *cis*- and *trans*-states (*λ* = 1), and we explore the role of the shape parameter *ϵ* (Fig. 3a). If *ϵ* = 1 (*cis* and *trans* have the same footprint on the membrane), *N*^*T*^ */N*_cz_ adopts a bell-shaped curve with a maximum at *µ* = −*E*_*b*_. This implies that for high bulk concentrations (*µ* close to zero), the system strongly prefers *cis* state over *trans*, inhibiting adhesion. Although such behavior is radically different from receptor-ligand bonding, where higher concentrations of tethers are expected to strengthen the adhesion, it is not unexpected since the enthalpy *density* per *cis*-tether is double that of a *trans*-tether - the two cholesterols of the *cis*-tether are inserted into the half of the membrane area of the *trans*-tether. The bell shaped curve is maintained until *ϵ* = 2, at which point the *trans* and the *cis*-tether contribute equally to the total free energy (*λ* = 1). Consequently, there are the same number of *cis* and *trans* tethers in the *µ* = 0 limit. In the regime *ϵ* > 2, the system behaves similarly to the receptor-ligand systems, with increasing tether concentration resulting in the increase of adhesion-promoting *trans*-tethers and therefore strengthening of adhesion, albeit being controlled in a very different manner. If the cost for making the *trans* bond increases (decreasing *λ*) the *trans* density rapidly decreases for all *µ* (Fig. 3b). However, binding can still be achieved by increasing *ϵ*.

This leads to the striking conclusion that the difference in the projected areas between states alone can drive contact formation due to collective entropic considerations. Even more strikingly, this remains true even if the adsorption affinity of the non-adhesive *cis*-state is larger than that of the adhesion-promoting *trans*-state, or, in other words, the *cis* state is on the single-molecule level energetically more favourable. It is important to note that the shape-factor *E* enters through entropic considerations in eq. 5. Its effect on contact-formation increases with bulk tether density because only in the crowded regime, where *cis* and *trans* states compete for adsorption area, do their projected area sizes become important. This can also clearly be seen in Fig. 3, where the low tether concentration regime does not depend on *ϵ*. Adhesion is therefore driven purely by collective effects and not by individual tethers. This indeed may be particularly relevant to the DNA-tethers where *λ* is indeed expected to be somewhat smaller than one, and *ϵ* a little bigger than two.

Finally, we inspect the tendency of the contact-zone growth by visualizing the change in the system’s energy Ω with the change of the contact zone size *N*_cz_ - captured by the so-called spreading pressure ∂Ω*/*∂*N*_cz_ (Fig. 3c). The observed negative spreading pressure corresponds to the tendency of the contact zone to spread, and higher tendencies for contact-zone spreading are highly correlated with higher densities of the *trans* bonds (compare Figs. 3b and 3c). We therefore expect *trans*-mediated adhesion to drive the spreading of the contact zone. However, unlike in the ligand-receptor binding, spreading tendency does not depend on the current size of the contact zone. The consequences of this size insensitivity would be very different response to forcefully induced de-adhesion.

### Membrane contact-formation assay

#### Experimental system

To quantify tether-mediated membrane contact formation, we let giant unilamellar vesicles (GUVs) interact with DNA-tether decorated SLBs. GUVs are introduced into a chamber containing an SLB already incubated with DNA-tethers. Due to the high adsorption affinity, tethers incorporate into the SLB as well as GUV membrane, probably both in *cis* (‘U’) configuration. GUV sedimentation is followed by formation of close contacts with the SLB. The GUV/SLB interaction is recorded using reflection interference contrast microscopy (RICM). It is seen that often the GUVs form an expanding contact zone with the SLB, where the two membranes are closely apposed. In this zone, referred to as the adhesion domain in adhesion literature, the two membranes are held together by DNA-tethers in *trans* (‘I’) configurations (Fig. 2c). In the absence of tethers, no formation of adhesion domains and no widening of the contact zone is observed. If tethers are present, in all cases we obtain a circular adhesion zone, leading us to infer that the adhesion proceeded through the formation of a radially growing adhesion patch [9]. This patch is formed by the transitioning of adsorbed *cis* tethers to the *trans* configuration through the re-insertion of one of the cholesterols into the apposing membrane. Alternatively, tethers can be adsorbed in *trans* configuration directly from the bulk solution. Both processes lead to adhesion (Fig.1a) and result in identical end-states that are indistinguishable in an equilibrium analysis.

#### Quantification of contacts

Visual inspection of RICM images reveals that in the same sample, some vesicles simply hover close to the SLB (non-adhered), some adhere exhibiting many fringes (strong adhesion), some adhere with just a few fringes visible (very strong adhesion) and few have no fringes at all (complete adhesion). Fig. 4a illustrates the different adhesion states that are encountered. The defined visual categories can be attributed objectively by defining an “adhesion parameter” *f*, which is given by the ratio of the diameter of the adhesion patch and the diameter of the GUV. *f* for each category is: *No Adhesion* 0 < *f* < 0.15, *Strong* 0.15 < *f* < 0.5, *Very Strong* 0.5 < *f* < 0.75, *Complete* 0.75 < *f* < 1.

**Figure 4.**
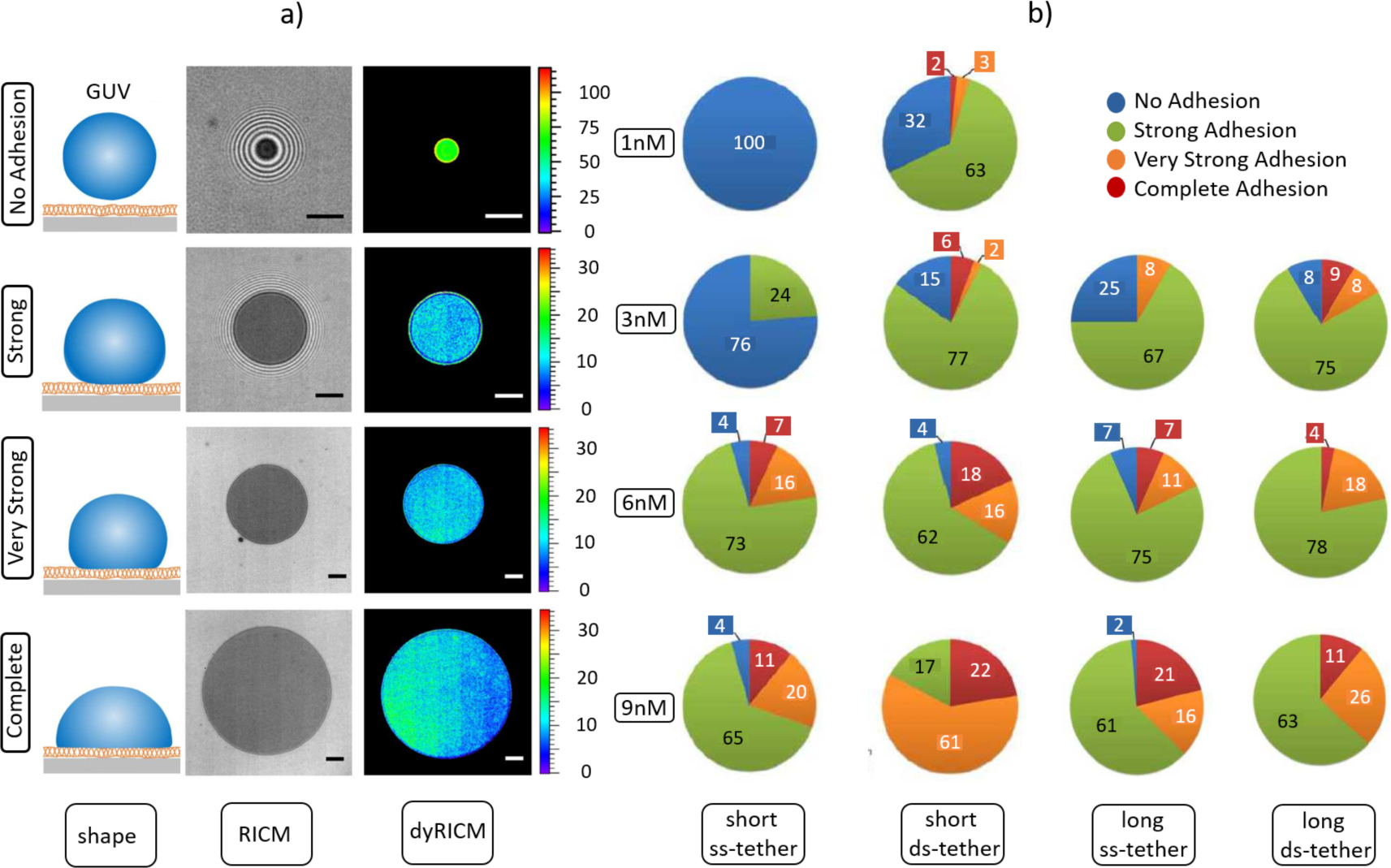
a) Different adhesion states identified on the basis of the ratio between the adhesion disc diameter and vesicle diameter. Note that the *Strong* case, where the energy can be estimated, shows that all the vesicles deemed to be in this state have the same adhesion energy, about 0.02*µJ/m*^2^, irrespective of the binder properties. The scale bar corresponds to 5 *µ*m. b) Comparison of the adhesion states between the ss-tethers and ds-tethers of two different lengths as a function of concentration. 1^*st*^ (2^*nd*^) column shows ss- (ds-tether) of length 11nm, while 3^*rd*^ (4^*th*^) column shows ss- (ds-tether) of length 19nm. Long ss-tethers, on average, have larger adhesion efficiency than short ss-tethers. On the other hand, short ds-tethers are on average more efficient than long ds-tethers.

Though we attributed names to the adhesion states based on morphology, only the so-called *Strong* adhesion state can be quantified using the Bruinsma construction which uses the Young-Dupre law with an *ad-hoc* line tension [19, 22] to infer adhesion energy density. We calculated the free energy in this state and find that all vesicles, irrespective of tether type, identified to be in the *Strong* adhesion state have an adhesion energy ranging from about 0.01 to 0.04 *µ*J/m^2^, and the final membrane tension of about 0.1 to 0.5 mN/m. Importantly, all the vesicles are equally strongly adhered independently of the tether type - short-ss, long-ss, short-ds and long-ds - implying that the classification by visual inspection is consistent.

#### Concentration dependence

We furthermore, characterize the extent of contact formation by counting the number of GUVs in each of these adhesion states, which depends strongly on the bulk concentration, as can be seen in Fig. 4b. Specifically, increasing the bulk concentration consistently increases the average contact zone size, independently of the tether type. This can be understood in terms of the balance between the spreading pressure, defined as ∂Ω*/*∂*N*_cz_ (Fig. 3c), which gives the tendency of a contact zone to spread. Comparing Figs. 3b and 3c, shows that for *ϵ* ≥ 2 the spreading pressure is a monotonous function of the chemical potential, and increases with the bulk density. In agreement with the observations of the increased frequency of larger contact zones at higher bulk concentrations of tethers (Fig. 4b), the result is the increased tendency for adhered contact zones to grow until the spreading pressure is not counteracted by the cost in the membrane bending energy.

To understand the dispersion in the adhesion states, we first need to realize that the assumption of the coupling to the reservoir of a constant chemcal potential makes the spreading pressure independent of the contact zone size. Therefore, smaller contact zones have the same tendency to grow as larger contact zones which are only limited by the GUV deformation energy that becomes large in the limit of a spherical cap. This means that the growth is arrested simultaneously for all vesicles. The dispersion over the various adhesion states not only depends on the variations in the mechanical properties of GUVs (mainly size and tension [9]) but also on the sedimentation time of each vesicle, which is a very different situation compared to the ligand-receptor driven adhesion.

#### Tether length and flexibility

Besides being controlled by the tether concentration in the solution, the observed adhesion state significantly depends on the tether type and its length. Interestingly, ss-tethers and ds-tethers exhibit qualitatively different behavior when their length is changed (Fig. 4b). It can be seen in Fig. 4b that the long ss-tether is, on average, more efficient at inducing adhesion than the short ss-tether, behaving in this context as a typical receptor. This should stem from the flexible nature of the construct for which the difference in projected area and the entropic cost of transitioning from the *cis* to the *trans* state would not be very significant (*ϵ* ≃ 2 and *λ* ≃ 1). As such, the ss-tether is behaving similar to a spring of constant stiffness, with the shorter spring costing more energy to deform the tether-membrane system and form a bond, resulting in short ss-tether being less effective at adhesion then the long ss-tether, as observed.

In contrast to the ss-tether behavior, short ds-tether is on average more efficient than the long ds-tether (Fig. 4b). Note that the ds-tether can be considered as composed of rigid rods connected by flexible joints, so the longer the ds-tether rods, the larger its projected area (*ϵ* > 2), and the higher is the entropic cost of being in the *trans* state relative to being in the *cis* state (*λ* < 1). We suggest that this is the reason for the observed adhesion efficiency of the shorter ds-tether compared to the longer ds-tether.

#### Gap-width in the contact zone

We also exploit the fact that RICM is almost the unique tool to measure the distance between two interacting membranes. As could be expected, the inter-membrane distance (*h*) increases with increasing tether length (Fig. 5a). However, *h* is independent of the bulk concentration of tethers, suggesting that a relatively high density of *trans* tethers is obtained in all cases. This is furthermore confirmed by the values of membrane roughness (defined as the spatial variation in *h*), which also adopts values independent of the tether concentrations. *h* depends weakly on the tether type (ss or ds), the gap being wider for ds-tethers.

**Figure 5.**
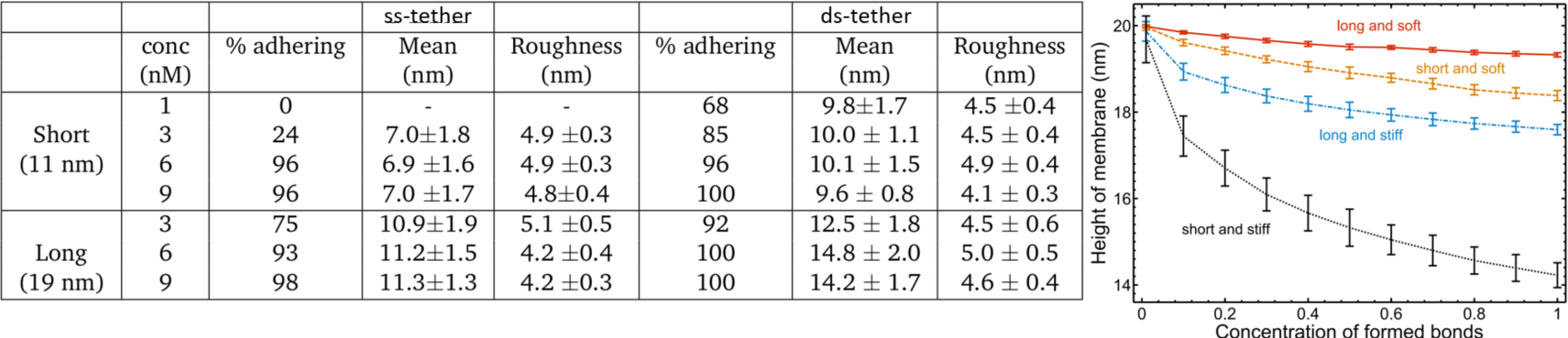
a) Table showing the percentage of adhered vesicles (all adhered states including week adhesion), the mean height and the roughness for all the cases studied. About 70 vesicles were recorded in each case, the heights were calculated on adhered vesicles, roughness is calculated as the standard deviation of height within the adhesion zone of each vesicle, the errors reported are the standard deviation of the entire population for each case. b) MC simulations. In the adhered state, the inter-membrane distance and roughness are uniquely determined by the tether length and flexibility, in a manner that is universally applicable. Adhesion mediated by longer tethers has larger inter-membrane distance. Stiffness of the tether increases the sensitivity of the inter-membrane distance and roughness on tether length.

These findings are supported by our theoretical calculations of the mean membrane height and roughens using the well-studied formalism for receptor-ligand cell adhesion [26]. Here, *trans* configurations are modeled as elastic springs pinning the membrane. By varying their density, length and stiffness we reproduce the qualitative behaviour observed in experiments (Fig. 5b), as long as the rigidity of the tether is not very high. The roughness (presented by error bar) is calculated as the true variation in membrane height averaged over the surface area. The height and the roughness of the membrane both decrease with density and independently of tether type, and both saturate to a constant value when the surface coverage of tethers in *trans* configurations exceeds a threshold density. However, the density dependent changes in roughness are very weak, and with the experimental resolution of few nanometers, they are not likely to be observed. Interestingly, the membrane height is sensitive to both the rest length of the tether and its stiffness. For a given stiffness, *h* is of course larger for longer tethers. For a given length, the softer tethers tend to support wider gaps. Comparing with experiments, we conclude that a jointed-rod configuration (ds-tethers) may in fact appear softer than a polymer-like configuration (ss-tethers), due to the very flexible joint. Over all, the gap-width can be regulated by adjusting either the length or the overall effective stiffness of the tethers.

## Conclusion

The system presented here introduces tethers that are soluble, rather than those that are permanently membrane bound. From a statistical physics point of view, this ensures that the chemical potential is maintained constant on all the surfaces and the bulk. In cellulo, this would correspond to a free exchange between the contact-forming membranes and the cytoplasm. While, this changes the statistical ensemble that needs to be used in the calculation and increases the importance of the entropy term, this alone does not introduce a large qualitative difference compared to the case where tethers are membrane-bound. However, if in addition, there is a conformation change upon formation of membrane bridges, the consequences are far-reaching. The geometric parameter associated with change in conformation, gets coupled to the surface density of the tethers on both membranes. The cytosolic concentration of tethers then becomes a potent tool to regulate membrane-bridging and contact formation. This is even more important if the *trans* configuration is costlier than the *cis* configuration - that is to say, if individual bonds are, in fact, energetically unfavorable. In this case, there is no binding expected without the cooperative effect induced by the geometrical parameter. In the DNA-tether system presented here, the *trans* configuration may in fact be costlier, since it involves pulling and confining a polymer, the tether, to a given length corresponding to the inter-membrane separation. This mechanism can of course work only if the tethers are soluble and can exist both in the cytosol and on the membranes.

In traditionally studied ligand-receptor systems, tether length and flexibility are expected to be important control parameters. Here, double or single stranded DNA was used to mimic soft or stiff tethers. Intriguingly, it turns out that the double-stranded DNA behaves as though it was the softer system - due to the fact that there are hinges around which the rod-like ds-DNA fragments can freely rotate. This is a cautionary tale about how details of the molecular structure may dominate the mechanics. We have shown both experimentally and theoretically that length and flexibility become secondary, with the geometrical parameter, along with bulk concentration, being the paramount control variable. Nevertheless, we show that they can act as secondary parameters that determine the contact gap-size (intermembrane distance), which in turn determines size exclusion - that is to say, it determines which molecules, whether soluble or membrane bound, can enter the contact-site and which cannot.

Here we presented prediction and in-vitro experimental verification of how this novel class of tethers may behave. The question arises whether this possible mechanism is exploited by nature. Even though identification of tethers and elucidating their dynamic structural parameters is very much in its infancy, there is at least one candidate - the Munc13 complex - which may have similar geometry [1]. The Munc13-Munc18 complex plays an important role in the capture of synaptic vesicles, forming a tripartite system close to the one described here. Even though the presence of three parts can be expected to modify the kinetics of contact formation, the essential steady-state/equilibrium features are already captured in our quantitative model.

The possibility of having tethers that can be both soluble and membrane-inserted provides the cell with a powerful toolbox that can be exploited for controlling intra-cellular adhesion-like processes. The most intriguing aspect is the revelation of conformation as a key parameter in inducing stable contacts, purely due to thermodynamic reasons totally separate from conformation induced affinity changes of single molecules. In particular, the cooperative nature of the process dominates over the single-tether level properties, and leads to contact formation even if it is energetically forbidden at a single molecular level. We show here that combination of solubility and size mismatch between *cis* and *trans* conformations, leads to totally new ways of controlling membrane-membrane contact formation. Such knowledge will help structural biologists to propose molecular mechanisms that were hitherto not considered to be within the realm of possibility.

## Acknowledgments

K.S. and A.-S.S. thank the joint German Science Foundation and the French National Research Agency project SM 289/8-1, AOBJ: 652939/ANR-18-CE92-0033(microCJ). J.A.J. and A.-S.S. thank European Research Council Starting Grant MembranesAct 337283. J.A.J. was in part supported by Croatian Science Foundation project DOK-2018-01-9055. L.L. was further supported by the China Scholarship Council (CSC, File No. 201806185038). MAK and KS acknowledge AMIDEX grant AFFINITY.

## Author contributions

K.S. and A.-S.S. directed and supervised the project. F.T. was in charge of designing, characterising and supervising the use of the tethers. M.A.K. and K.S developed the GUV-SLB system and performed the measurements and analysis. J.A.J. and A.-S.S provided theoretical modeling and applied it on experimental data. L.L. calculated the membrane height vs. concentration of *trans* states (Fig5b). K.S., A.-S.S., J.A.J. and M.A.K wrote the manuscript with contributions from all authors.

## Materials and methods

### Lipids and other reagents

1–stearoyl–2–oleoyl–sn–glycero–3–phosphocholine (SOPC), 1,2–distearoyl–sn–glycero–3–phospho ethanolamine–N–[amino(polyethylene glycol)–2000] (DSPE – PEG 2000), head-labeled 1,2– dioleoyl–sn–glycero–3–phosphoethanolamine–N–%[7–nitro–2–1,3–benzoxadiazol–4–yl] (NBD–PE), chain-labeled 1-oleoyl-2-6-[(7-nitro-2-1,3-benzoxadiazol-4-yl)amino]hexanoyl-sn-glycero-3-phosphoc (NBD-PC) and 1,2-dioleoyl-sn-glycero-3-phosphoethanolamine-N-(lissamine rhodamine B sulfonyl) (RhodamineB) were purchased from Avanti Polar Lipids (Alabaster, AL) and used without further purification. Dextrose (Glucose), BioXtra, ≥ 99.5% (GC) was purchased from Sigma Aldrich, France. The DNA-tethers were short custom-made single-stranded DNA-oligomers from (Eurogentech, Belgium).

### SLB, GUV formation

Supported lipid bilayers (SLBs) were prepared with a film balance (Nima, Coventry, UK) applying the Langmuir-Blodgett Langmuir-Schäfer technique [**?**]. The subphase was ultrapure water. SLBs consisted of pure SOPC in the proximal layer. The distal layer facing the buffer was formed by SOPC with 2 mol % DSPE-PEG 2000 and 1 mol % NBD-PC (or NBD-PE or RhodamineB). Transfer pressure was set to 20 mN/m. SLBs were constantly kept under water and used directly after preparation. Giant Unilamellar Vesicles (GUVs) were prepared using the well established electroswelling method. Briefly, 10 *µ*L of the lipid mixture (98 mol% SOPC and 2 mol % DSPE-PEG2000) dissolved in chloroform was spread on indium tin oxide coated glass slides (Sigma Aldrich, France). In order to ensure complete evaporation of chloroform the slides were desiccated under vacuum overnight. Two lipid-coated glass slides were mounted in a teflon chamber filled with 230 mOsmL^−1^ glucose solution, at a distance of 1 mm. An alternating voltage of 2 V at 10 Hz was applied for 2hrs which resulted in GUVs with an average diameter of around 20 - 30 *µ*m. For experiments, vesicles were immersed in PBS buffer of 270 mOsmL^−1^. The difference in osmolarity between the inside and outside solutions resulted in vesicles exhibiting the necessary excess area to build up a contact zone with the substrate. All osmolarities were measured before each experiment with an osmometer (Osmomat 030, Gonotec GmbH, Berlin, Germany). 20 - 40 *µL* of the vesicle solution was added into the experimental chamber filled with PBS buffer (volume 1 mL). Vesicles were allowed to sediment and achieve a steady adhesion state before the first measurement. The waiting time on an average was 30 mins. Observation chambers were filled completely with PBS and covered with a glass slide to avoid osmolarity changes due to evaporation. For height determination using RICM, the refractive indices of the vesicle solution and the outer buffer were measured for each experiment with an Abbé Refractometer (Kruss, Germany).

### Microscopy and data analysis

For quantification of DNA absorption, images were acquired using a confocal microscope (Leica Microsystems, Germany) fitted with a 63x long distance oil immersion objective. For all other experiments, images were acquired using an inverted microscope, Zeiss Axiovert 200, equipped with a EM-CCD camera (Andor, Ireland), a filter cube with crossed polarizers and a 63X Antiflex Plan-Neofluar oil objective (Carl Zeiss, Göttingen,Germany) having a numerical aperture of 1.25. In addition the objective had a built in lambda quarter plate. For reflection interference contrast microscopy (RICM), the green illumination was selected from the light emitted by a metal halogenide lamp (X-Cite, Exfo, Quebec, Canada) using an interference filter (546 ± 12 nm). The numerical aperture of illumination was set to approximately 0.5. Image sequences consisted of 50 consecutive frames with an individual exposure time of 100 ms for each observation set.The adhesion state of GUVs was probed using reflection interference contrast microscopy (RICM). Map of membrane-substrate distances from the measured interference patterns were constructed [14]. Briefly, the entire image (the vesicle as well as the background) was first corrected for anomalies arising from an inhomogeneous illumination. Next, the average background intensity (I_*bg*_) in the image was measured and the entire image was normalized with respect to I_*bg*_. Finally, the normalized intensities were used to find the corresponding height by inverting the intensity height relation [**?**]. For all quantitative evaluation, images were analyzed using self-written routines in IgorPro (Wavemetrix, Portland, OR).

## Appendix

### Derivation of the thermal equilibrium properties

Assuming thermal equilibrium, we demand

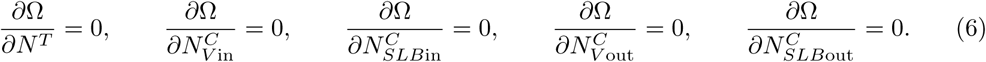

With the help of the Stirling approximation

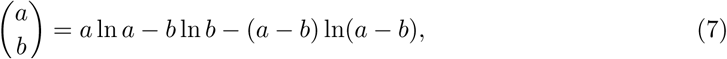

solving the coupled eqs. (6) for equilibrium numbers *N*^*T*^, 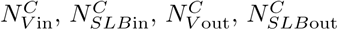 gives

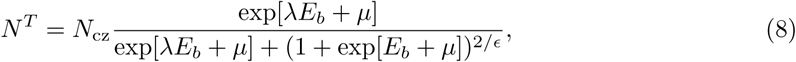

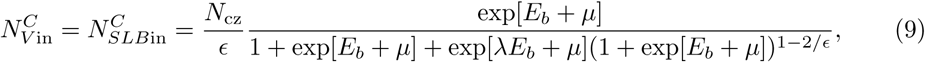

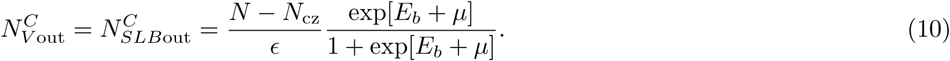

